# A Method for Sampling the Living Wood Microbiome

**DOI:** 10.1101/2023.11.14.567064

**Authors:** Wyatt Arnold, Jonathan Gewirtzman, Peter A. Raymond, Mark A. Bradford, Claire Butler, Jordan Peccia

**Author notes:** Contributed equally to this work (Wyatt Arnold, Jonathan Gewirtzman). Corresponding author: Jordan Peccia.

## Abstract

Efforts to characterize microbial life across diverse environments have progressed tremendously, yet the microbiome of Earth’s largest biomass reservoir—the wood of living trees—has been largely unexplored. Current understanding of the tree microbiome is largely confined to roots and leaves, with little attention given to the endophytic microbiome of wood, even though emergent studies have indicated this zone as a niche for unique taxa, of consequence for ecosystem health and global biogeochemical cycles. The lack of investigation derives partly from the physical recalcitrance of wood, which presents challenges during sampling, homogenization, and the extraction of nucleic acids. In response to these issues, we present an optimized method for processing wood for use in microbial analyses, from sampling through to downstream analyses. Using methane-cycling taxa as model endophytes, we assess losses in recovery during our method, and determine a limit-of-detection of approximately 500 cells per 100 mg of (dry) wood. For all six species evaluated—which represented several diverse taxa of hardwoods and softwoods—PCR inhibition proved minimal, and we expect this method to be applicable for a majority of tree species. The methods presented herein can facilitate future investigation into the wood microbiome and global microbial ecology of methane cycling.

## Introduction

Trees play a vital role in both ecological and economic systems, shaping their environments and supporting various industries, with forestry expected to eclipse a global valuation of $1 trillion in 2023 (*Forestry And Logging Global Market Report 2023*, 2023). In addition to their economic benefits, trees have an established and growing role in carbon capture and sequestration (Ni *et al*., 2016), and have historically served as a source of important pharmaceuticals (Pandey, 1998; Jones, 2011). As such, improving our understanding of the physiology and biogeochemical function of these conspicuous plants has clear economic and social benefit.

Just as the human microbiome directly affects a vast number of physiological and metabolic processes within an individual (Gilbert *et al*., 2018; Arnold *et al*., 2019), the known tree microbiome plays crucial roles in growth, reproduction, and response to environmental stress (Uroz *et al*., 2016; Terhonen *et al*., 2019; Mishra, Hättenschwiler and Yang, 2020). While microbes found in the rhizosphere (Habiyaremye *et al*., 2020), dermosphere (Jeffrey *et al*., 2021), and phyllosphere (Leveau, 2019) of trees are increasingly well characterized, comparatively little research has investigated the role of endophytic microbial taxa within the stems of trees, which constitutes the largest portion of their biomass (Baldrian, 2016), and comprise the largest pool of biomass on Earth (Bar-On, Phillips and Milo, 2018). Increasing evidence implicates tree stems as sites of notable microbial activity (Hacquard and Schadt, 2015), including methane-cycling metabolisms (Barba *et al*., 2019; Jeffrey *et al*., 2021; Putkinen *et al*., 2021), a fact which may have consequences for global greenhouse gas accounting (Vargas and Barba, 2019).

There are, however, practical difficulties involved in surveying the microbiome of living wood. For every step, from sample collection, to homogenization, to microbial nucleic acid extraction and downstream analyses, the physical resiliency of the material and its numerous inhibitory compounds present logistical challenges (Rachmayanti *et al*., 2009). While promising inroads into the study of wood-borne communities (Proença *et al*., 2017; Denman *et al*., 2018; Ren *et al*., 2019; Flanagan *et al*., 2021), and the methods for doing so (Verbylaite *et al*., 2010), have been made, a lack of a critically validated procedure for the extraction and amplification of microbial nucleic acids from living wood presents a significant barrier-to-entry for the field and complicates the comparison of studies which rely on different methodologies **(Table 1)**. Characterizing the limits of the analytical tools involved in the study of wood endophytes is essential for accurately interpreting the ecological and functional significance of these communities and understanding their role in forest ecosystems, tree health, and potential applications in biotechnology and conservation.

**Table 1:**
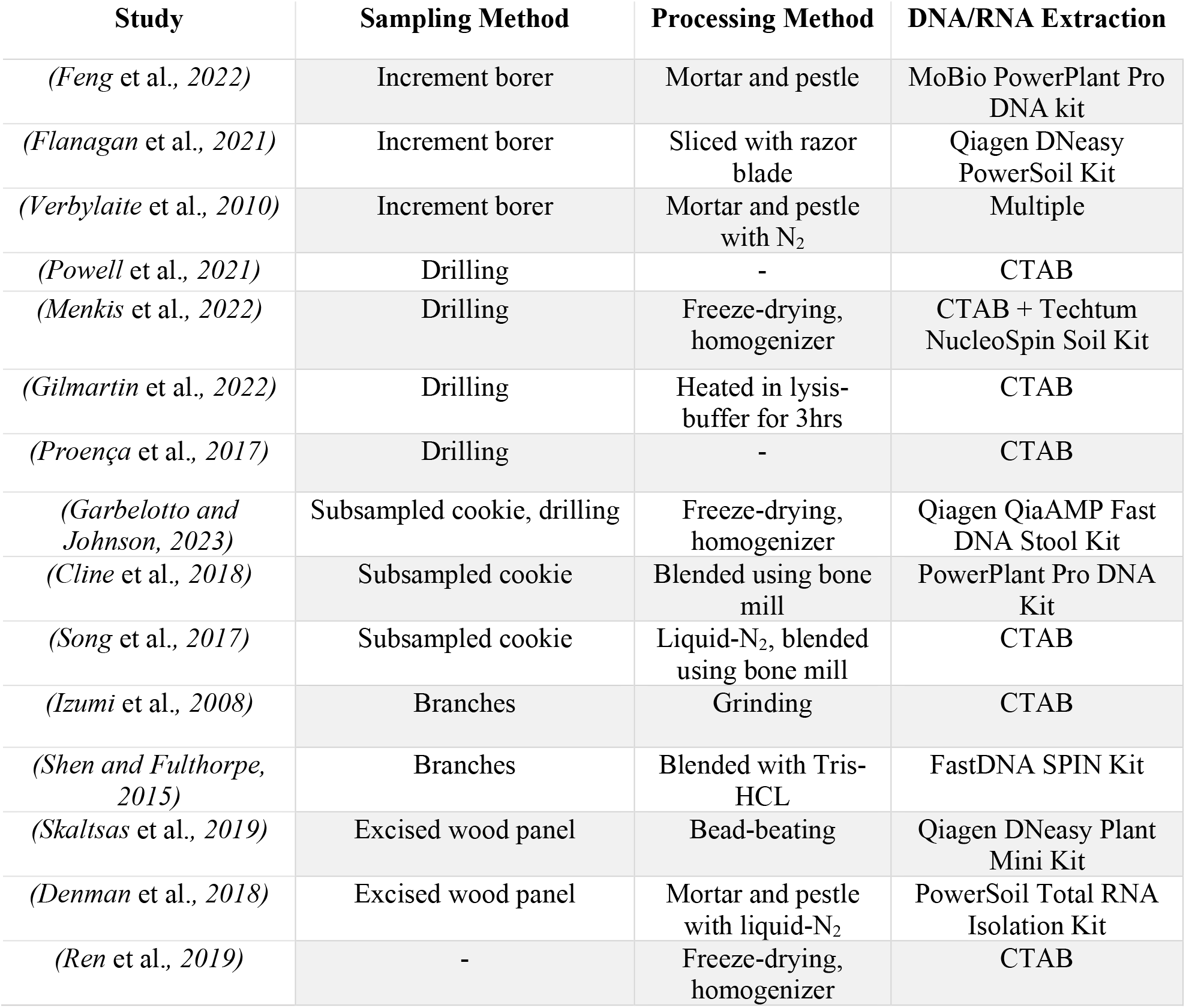
Overview (non-exhaustive) of publications wherein the endophytic wood microbiome was sampled. Listed for each publication are wood sampling methods, wood processing techniques, and nucleic acid extraction methods.

We present an approach—optimized over a year of trials—and accompanying evaluation thereof, for the extraction of microbial nucleic acids from the wood of living trees. Focusing on endophytic methane-cycling taxa as an analytical example, we describe our methodology developed to survey the constituents of the wood microbiome in Northeastern US temperate forests, and we identify resolved and continuing challenges faced when studying wood endophytes.

## Materials and Methods

### Sample Collection

Two sampling campaigns were conducted to source living tree wood samples for method development and validation. The first set of samples, collected in June 2022, were used for initial method development, and to inform assay development and DNA extraction kit selection. The second set of samples, collected in January 2023, were used for additional analyses of method performance. All samples were sourced from Yale-Myers Forest, a managed research and demonstration forest in Northeastern, CT, USA (41.9529° N, 72.1239° W). Soils at the site are primarily glacial-till, with defined organic and mineral horizons. The forest is classified in the Central Hardwood-Hemlock-Pine region (Westveld, 1956), and consists mostly of second-growth vegetation which developed following agricultural abandonment in the mid-19^th^ century. We selected a variety of dominant trees in the region to represent a wide range of species and wood traits, including both hardwoods and softwoods, deciduous and evergreen leaf strategies, and arbuscular and ectomycorrhizal associations. These selections were made under the assumption that both species and wood traits may present unique challenges for processing and downstream analyses. All trees sampled had a diameter at breast height (DBH, 1.37 m from the ground) of at least 51 cm (20 inches), were living at the time of sampling, and had no overt signs of disease or other stressors (drought, damage, etc.).

June 2022 samples were taken from 12 individual trees across four different species, two each from genera of angiosperms and gymnosperms: sugar maple (*Acer saccharum*) (n=3), black birch (*Betula lenta*) (n=2), Eastern hemlock (*Tsuga canadensis)* (n=4), and Eastern white pine (*Pinus strobus*) (n=3). January 2023 samples were taken from 12 individuals, with 3 trees cored for each of four species: Eastern hemlock, eastern white pine, red oak (*Quercus rubra*), and red maple (*Acer rubrum*). For January 2023 samples, duplicate cores were collected from each tree (n=24 total). Cores were taken using a 4-mm increment borer (Haglöf Sweden, Sweden), which was decontaminated in between samples by flame-sterilizing with ethanol. Immediately following collection, cores were sealed in sterile Al-foil and placed onto dry-ice for transport back to Yale University, after which they were stored at −80°C until further processing.

### Sample Processing

After collection, cores were subsampled into heartwood (“inner”) and sapwood (“outer”) sections to evaluate any differences between the two tissues during downstream analyses. Given known challenges of delineating a precise sapwood-heartwood boundary in certain species (Munster-Swendsen, 1987), we separated the inner 5 cm (as measured from the pith) and the outer 5 cm (as measured from the bark) of the cores. This length was chosen to yield sufficient material for extraction and so that processing occurred using equivalent sample volumes. All steps were performed on a chilled block (near 0 °C) according to sterile procedure (e.g. flame-sterilizing tools, decontaminating surfaces with bleach) within a laminar flow-hood to minimize environmental and inter-sample contamination.

Cores were then freeze-dried before being processed into a fine, homogenous powder to ensure a consistent sample for each tissue type and to transform the cores into an appropriate input for nucleic acid extraction (**Figure 1**). Freeze-drying reduces core moisture content to near zero, minimizing the effect that variable moisture content might have on grinding efficacy, limiting damage to nucleic acids from shearing that would otherwise be induced by ice crystal formation during subsequent steps, and allowing for standardization of results on a dry mass basis. During freeze-drying, individual samples were placed, frozen, into 15-mL centrifuge tubes outfitted with 0.22-μm filter caps (CELLTREAT Bio-Reaction Tubes, CELLTREAT, MA, USA) to enable gas exchange and were lyophilized for 72 hours using a flask freeze-drying system (Labconco, MO, USA). Water-loss was assessed through weighing, typically resulting in a ∼50% reduction in sample mass.

**Figure 1:**
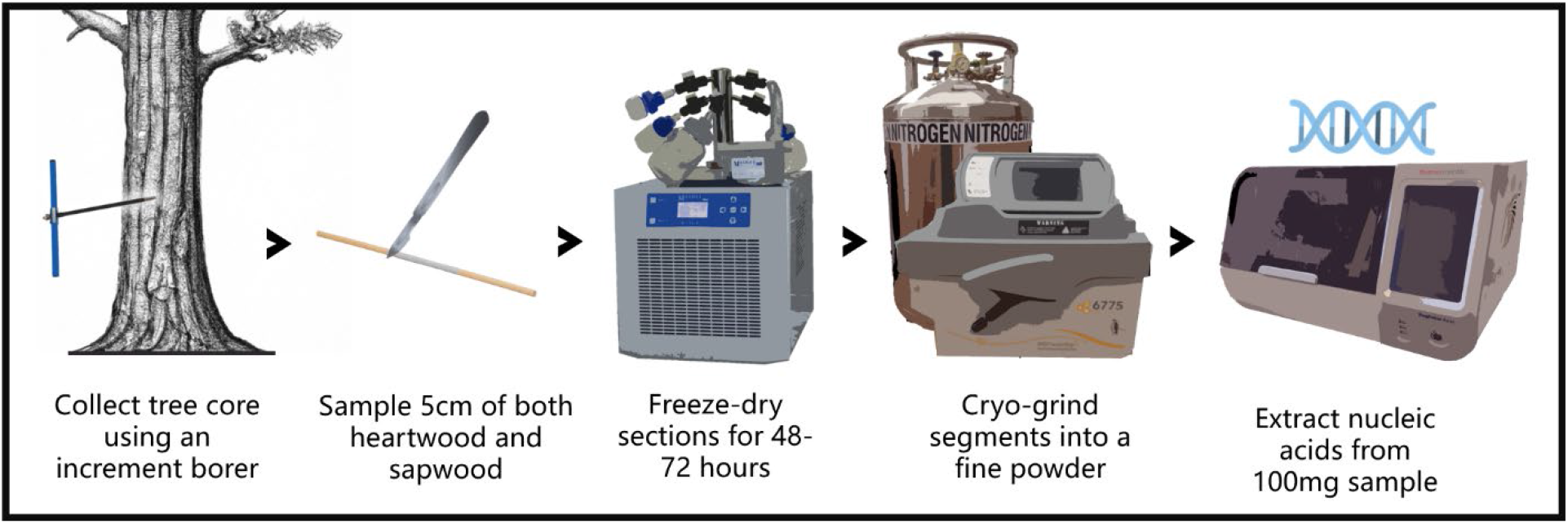
Overview of the collection and processing pipeline for extracting nucleic acids from wood samples.

Samples were then individually ground to a fine powder in a 6775 Freezer/Mill® Cryogenic Grinder (Spex, NJ, USA), which uses an electromagnetically driven impactor to grind individual samples within an easily decontaminated vial that is immersed in liquid N2 during the grinding process. The grinding cycle was as follows: pre-cool for 10 min, followed by 2 cycles of alternating running and cooling steps, each 2 min in duration. Grinding was performed at a rate of 10 cycles per second. The ground mass from a 5 cm segment ranged from around 300-600 mg depending on species and wood type.

While we explored grinding through alternative techniques, such as a Wiley® mill, mortar and pestle, ball mill, blender, and hand-grinding, we ultimately found these techniques unable to satisfy our criteria for sample handling. These criteria included the ability to conduct high-throughput processing, to quickly and completely decontaminate the grinding apparatus, to create a homogenous, fine (no visible fibers), and consistent powder, and to impose minimal damage to the sample’s nucleic acids through heat generation or mechanical degradation.

### Spiking Experiments

To 1.) determine a suitable nucleic acid extraction kit, 2.) quantify the effects of sample handling procedures, and 3.) evaluate the dependency of recovery efficiency on tree species and tissue type, a proportion of samples was spiked with sufficient nucleic acids to yield a positive analytical signal (**Table 2**). As quantitative evaluation was performed via digital droplet PCR (ddPCR), spiking material included both synthetic dsDNA oligonucleotides and whole cells.

**Table 2:** Overview of spiking experiments for method development and evaluation. The number in parentheses next to the spike type represents the number of targets spiked (e.g. mcrA and pmoA [2]).

| Samples | Use | Spiked prior to: | Freeze-Drying | Cryo-Grinding | Extraction |
| --- | --- | --- | --- | --- | --- |
| *Sterile Dowels* | LoD analysis, inhibition testing, DNA sequencing | Sample Set 1 (n=3) | gBlock (3) |  |  |
|  |  | Sample Set 2 (n=3) | Culture (2) |  |  |
|  |  | Sample Set 3 (n=3) |  | gBlock (3) |  |
|  |  | Sample Set 4 (n=3) |  | Culture (2) |  |
|  |  | Sample Set 5 (n=3) |  |  | gBlock (3) |
|  |  | Sample Set 6 (n=3) |  |  | Culture (2) |
|  |  | Control (n=3) |  |  |  |
| *January, 2023* |  | Field Spike (n=24) |  |  | gBlock (3) |
|  |  | Field Control (n=24) |  |  |  |
| *June, 2022* | Kit selection | Field Spike (n=12) |  | gBlock (2) |  |
|  |  | Field Control (n=12) |  |  |  |

For three different targets, synthetic dsDNA (gBlocks) regions were produced, (Integrated DNA Technologies, IA, USA) ranging from 350 to 500 bp in length. Two of the targets were derived from the genome of *Methylococcus capsulatus*, one a fragment from a region encoding the pmoA gene and the other from a region encoding the mmoX gene (both methane monooxygenase genes involved in methane oxidation). The third target was derived from the genome of *Methanococcus voltae*, from a region encoding the mcrA gene (alpha subunit of methyl-coenzyme M reductase, involved in methanogenesis).

As an additional metric to evaluate sample recovery, cultured cells were also spiked onto a subset of tree cores. *Methanococcus maripaludis* (ATCC 43000), a methanogenic archaeon containing the mcrA gene, was grown on liquid Methanogenium medium (ATCC 1439) in sealed mesocosms under an 80% H_2_, 20% CO_2_ atmosphere (pressurized to 20 psi). Mesocosms were kept at 37°C on a shaker plate, with the atmosphere replaced weekly. *Methylococcus capsulatus* (ATCC 33009), a methanotrophic bacterium with both pmoA and mmoX genes, was grown on liquid NMS media (ATCC 1306) in sealed mesocosms under a 50% CH_4_, 50% sterile air atmosphere. Mesocosms were kept at 37°C on a shaker plate, with the atmosphere replaced weekly. This choice of taxa—which contained the mcrA, pmoA, and mmoX genes—ensured the same ddPCR methods could be used for quantification across both spike types.

To assess loss due to each processing step, 6.35-mm diameter Baltic birch (*Betula spp.*) dowels were used as the basis for a controlled spiking experiment. This material was selected over field-collected samples as the field samples often displayed structural variability which was likely to confound results. Dowels, in triplicate sets, were spiked at each point in the processing and extraction pipeline (**Table 2**) to assess loss at each sequential step. For the samples spiked prior to cryo-grinding, either 50 μl of each of the two cultures, directly sampled from mesocosms, or 50 μl of three 0.1 pg μl^-1^ gBlock solutions (mcrA= 166,619 copies μl^-1^, pmoA=129,801 copies μl^-1^, mmoX= 247,301 copies μl^-1^), were pipetted across the surface of 3 sterile dowels and allowed to soak in. For the samples spiked after cryogrinding, a comparable mass of the same 3 gBlock targets or culture was pipetted directly into the homogenized powder, vortexing briefly to ensure even dispersal. This amount of spiking material was chosen to ensure that, when analyzed via ddPCR, the spiked samples would fall within the linear range of the instrument.

Additionally, to understand potential dependency of extraction efficiency on species, wood type and tissue (heartwood vs. sapwood), triplicate, spiked extractions were performed using material collected from the four species sampled in January 2023. To obtain a consistent sample for analytical replicates, the field-collected cores were combined into single sets for each species, producing one bulked heartwood and one bulked sapwood sample per species for downstream processing and analysis. Bulked samples were individually freeze-dried and cryo-ground. Immediately prior to extraction, triplicate subsamples of the homogenized products received approximately 25 μl of each (0.1 pg μl^-1^) of the 3 gBlock targets (mcrA, mmoX, and pmoA).

Lastly, using the 24 samples (12 species x heart/sapwood) produced during the June 2022 sampling campaign, half were spiked with mcrA and pmoA gBlocks after lyophilization for use in a comparison of nucleic acid extraction kits. gBlocks were selected over cultured cells as we could better quantify the expected number of gene copies, and we expected minimal differences in lysis across kits given they both use similar lysis (detergent and bead-beating) methods.

### DNA Extractions

For samples collected in June 2022 and January 2023, DNA was extracted from 100 mg of cryo-ground sample using a MagMAX™ Microbiome Ultra Nucleic Acid Isolation Kit in concert with a KingFisher Apex automated nucleic acid extraction system (Thermo Fisher Scientific, MA, USA). Prior to extraction, samples (100 mg lyophilized powder, supplied beads, and 800 µL lysis buffer) were bead-beat for 10 min at maximum speed on a Vortex-Genie (Scientific Industries, NY, USA). The standard 96-well extraction protocol was followed for the kit using the manufacturer’s program (KingFisher Apex MagMAX_Microbiome_Soil_Liquid_Buccal_v1.kfx). Samples were ultimately eluted into 75 μl of supplied elution buffer.

To compare the recovery of nucleic acids from the above protocol with an extraction kit made for plant materials, duplicate extractions were performed on the June 2022 samples using the MagMAX™ Plant DNA Isolation Kit (Thermo Fisher Scientific, MA, USA) in concert with a KingFisher Apex automated nucleic acid extraction system. Again, 100 mg of sample was used for extraction, samples were bead-beat for 10 min on a vortexer prior to extraction with polyvinylpyrrolidone added to Lysis Buffer A at a 2% (w/v) concentration per manufacturer recommendations for woody samples. The manufacturer’s standard extraction protocol and program were (KingFisher Apex MagMAX_Plant_DNA_v1.kfx) followed, with the nucleic acids eluted into 75 μl of supplied elution buffer.

### Inhibition Testing

Plant samples, especially woody, lignified samples, are known to contain compounds inhibitory to downstream enzymatic applications, such as polyphenols, which are abundant in heartwood of many tree species (Scalbert, 1992). Due to similar sizes and charges, these compounds often coextract with nucleic acids. As such, inhibition of ddPCR was evaluated in two different ways. First, a dilution series was performed on the spiked field samples sourced in January 2023 to investigate inhibition across species and tissue-type. Sample eluents were diluted, in triplicate, at 1:5, 1:25, and 1:125 ratios with IDTE buffer (Integrated DNA Technologies, IA, USA). The probe-based mcrA ddPCR assay (described below) was used as the method of quantification for these tests. Second, for the June 2022 samples, a Zymo OneStep PCR Inhibitor Removal Kit (Zymo Research, CA, USA) was used to purify the extracted nucleic acids of all samples (n=24). This kit is specifically designed for the removal of compounds such as polyphenols and tannins from samples. ddPCR, using EvaGreen chemistry, and targeting the pmoA gene, was run both before and after purifying the samples.

### ddPCR Techniques and Assay Development

All quantification was performed via ddPCR using a QX200 Droplet Reader, along with a QX200 Droplet Generator (Bio-Rad Laboratories, CA, USA). Reaction mixes and thermocycling schedules are listed in **SI Table 1**. Assays targeting the pmoA, mmoX, and, initially, mcrA genes relied on EvaGreen chemistry, a dsDNA intercalating dye, and used preexisting primer sets from the literature (Fuse *et al*., 1998; Bourne, McDonald and Murrell, 2001; Luton *et al*., 2002). Early testing revealed, however, that the mcrA assay relying on EvaGreen chemistry was prone to a low, but persistent rate of false positives (**Figure 2**) in wood. Moreover, the rate and degree of false positives appeared dependent on species and tissue, further complicating usage of the primer set (**SI Figure 1**). While the assay would be sufficient for high abundance (e.g., soil) samples, we expect DNA eluants from healthy wood samples to harbor low (1-50 copies) abundances of target microbial sequences, and thus we sought an improved method with greater sensitivity and reliability. As probe-based assays are known to convey these improvements, we opted to develop a degenerate probe-based assay for the mcrA region (**SI Figure 2**).

**Figure 2:**
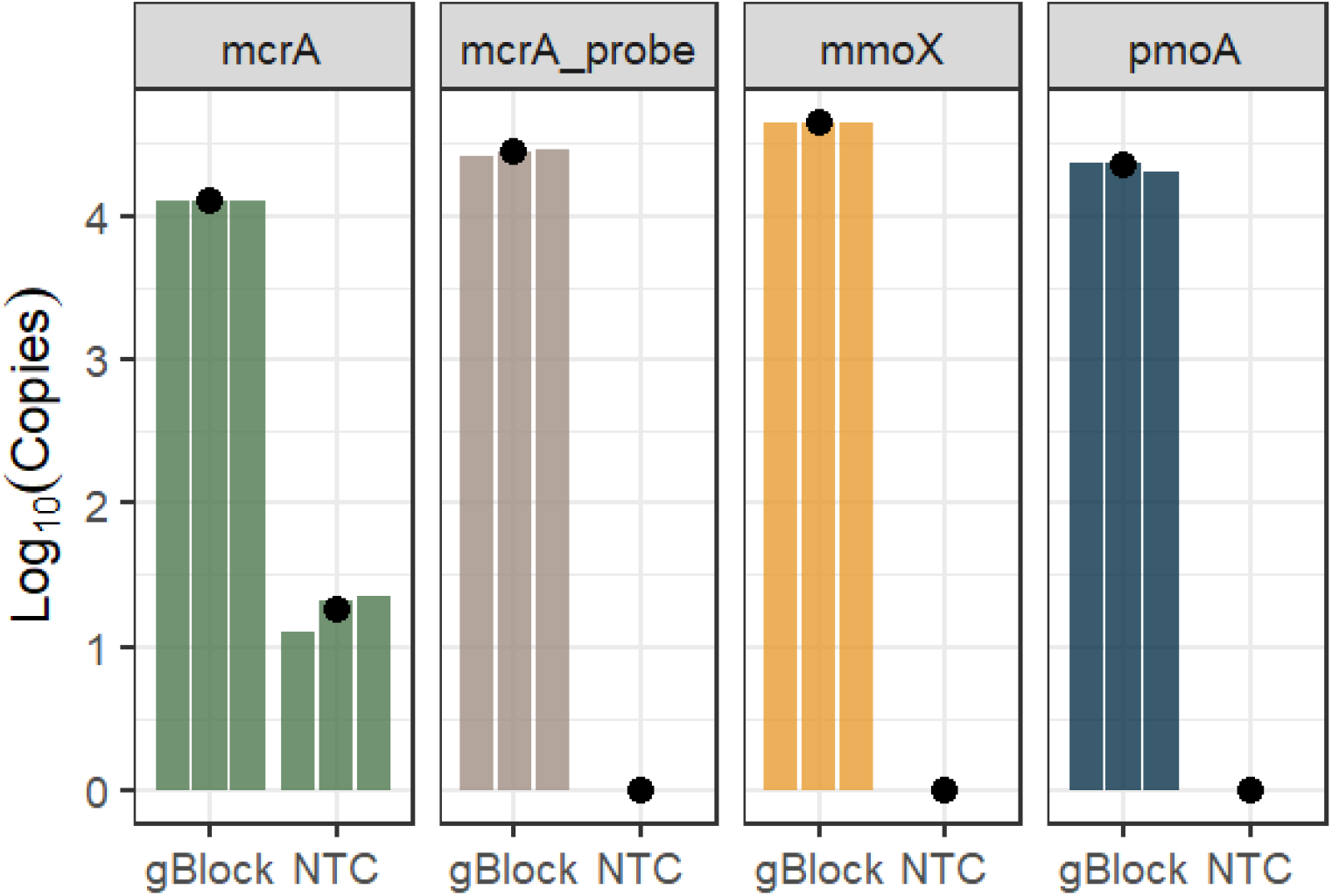
Triplicate positive (gBlocks, 10^4^ copies expected) and no template controls (NTC) for ddPCR assays used for quantification of methanogenic (mcrA, mcrA_probe) and methanotrophic (mmoX, pmoA) taxa in extracted DNA. Average in black. All assays are EvaGreen-based except for the mcrA_probe, which is a newly designed degenerate probe-based assay (**SI Table 1**).

Primer and probe development was performed in Geneious Prime (Dotmatics, MA, USA) using sequences sourced from the NCBI database (Sayers *et al*., 2022). In total, 75 RefSeq sequences for the coenzyme-B sulfoethylthiotransferase subunit alpha (mcrA) gene were aligned using Clustal Omega (Sievers *et al*., 2011). Two sets of degenerate primers and probes were then designed based-off of the consensus sequence (75% threshold) using the primer3 algorithm (Untergasser *et al*., 2012). Both sets produced an amplicon that was 118bp (**SI Table 1**) in length. To investigate the effect of fluorophore choice, one probe was synthesized with FAM as the fluorophore, and the other with HEX. Primers were synthesized at the Keck Oligonucleotide Synthesis facility at Yale University. Probes were synthesized by IDT using their PrimeTime chemistry. For both the FAM and HEX probes, quenching was accomplished via Iowa Black® FQ with an internal ZEN quencher. Primer and probe sequences, along with thermocycling conditions and reagent mixes are listed in **SI Table 1**.

### Amplicon Sequencing and Bioinformatics

DNA, extracted from cores sourced from three additional Red Maple trees at Yale-Myers Forest, was sent to The University of Minnesota Genomics Center (UMGC) for amplicon sequencing. Universal primers (515F/806R) targeting the V4 region of the 16S rRNA gene were selected. In order to limit amplification of plastid and mitochondrial DNA, PNA (mPNA and pPNA) blockers (PNA Bio, Newbury Park, CA, USA) were added during library construction. Libraries were dual-indexed, and subsequently sequenced on a 2 x 300 bp paired-end flow cell using the Illumina MiSeq platform (Illumina, San Diego, CA).

Pre-processing of sequence data was accomplished using the *dada2* package v1.18 (Callahan *et al*., 2016), with computation performed using Nephele (Weber *et al*., 2018). Subsequent data processing was then performed in R v.4.0.4 (R Core Team) using the *phyloseq* package v.1.34.0 (McMurdie and Holmes, 2013). Prior to rarefying at 8,000 reads per sample, all chloroplast and mitochondrial sequences were filtered out.

## Results and Discussion

### Extraction Method Selection and Inhibition

Overall, the MagMAX Plant and MagMAX Microbiome kits performed similarly, with no significant difference in the recovery of the gBlocks across the twelve spiked samples (paired t-test, p>0.05) (**Figure 3**). While both kits appeared sufficient for extraction of nucleic acids from wood, the remainder of experiments were conducted using the more generalist Microbiome kit, as it simultaneously recovers both DNA and RNA, can accommodate other sample types of interest (e.g. soil), and has been validated as a kit-of-choice in amplicon sequencing (Shaffer *et al*., 2022).

**Figure 3:**
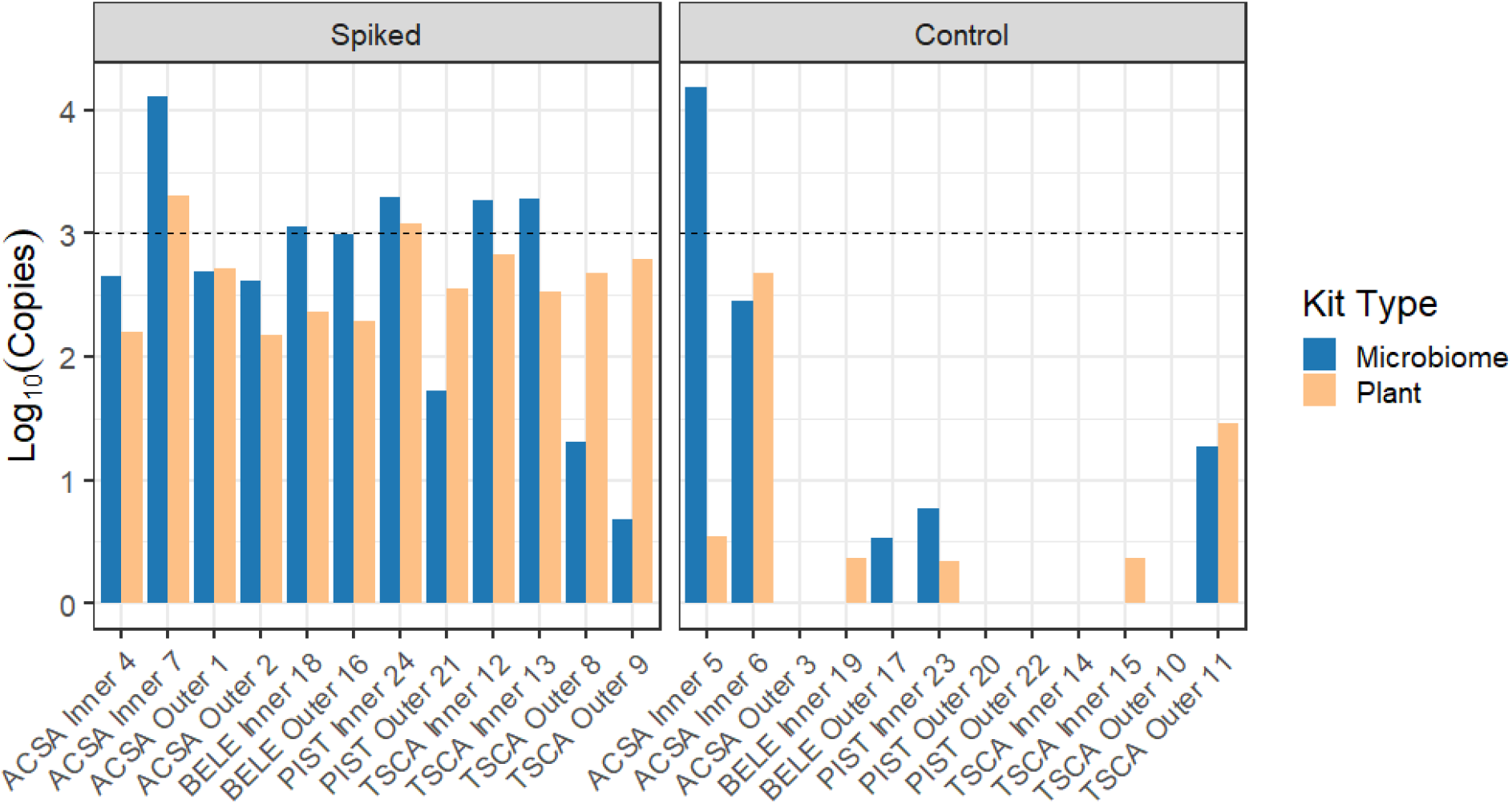
Comparison of 24 tree wood samples, 12 spiked and 12 control (no added gBlocks), extracted using two different kits and quantified using the mcrA probe-based assay. Samples were spiked with a mass of gBlocks expected to yield a positive signal of approximately 10^3^ copies (dashed line). Across the 12 spiked samples, there was no difference in recovery for the two kits (p=0.97). Each x-axis tick mark label represents an individual tree sample (ACSA: *Acer saccharum*, BELE: *Betula lenta*, PIST: *Pinus strobus*, TSCA: *Tsuga canadensis*), with the number serving as just a unique identifier.

PCR inhibition was not apparent for any of the sample types tested, and as such showed little dependency on tree species, tissue type, or clade. The average slope of the dilution series, produced using the four species collected in Jan 2023, which were spiked with gBlocks and extracted with the MagMAX Microbiome kit, was 0.98, with an R^2^>0.99 and a p-value<0.001 (**Figure 4**). The linearity of these dilution series, and proximity of the slopes to 1, indicates that the expected and observed dilution values matched well across a wide range of concentrations, implying minimal inhibition. Moreover, this result was further corroborated by testing with a Zymo OneStep Inhibitor Removal Kit, wherein no significant difference (paired t-test, p>0.05) was observed in a comparison of treated and untreated samples.

**Figure 4:**
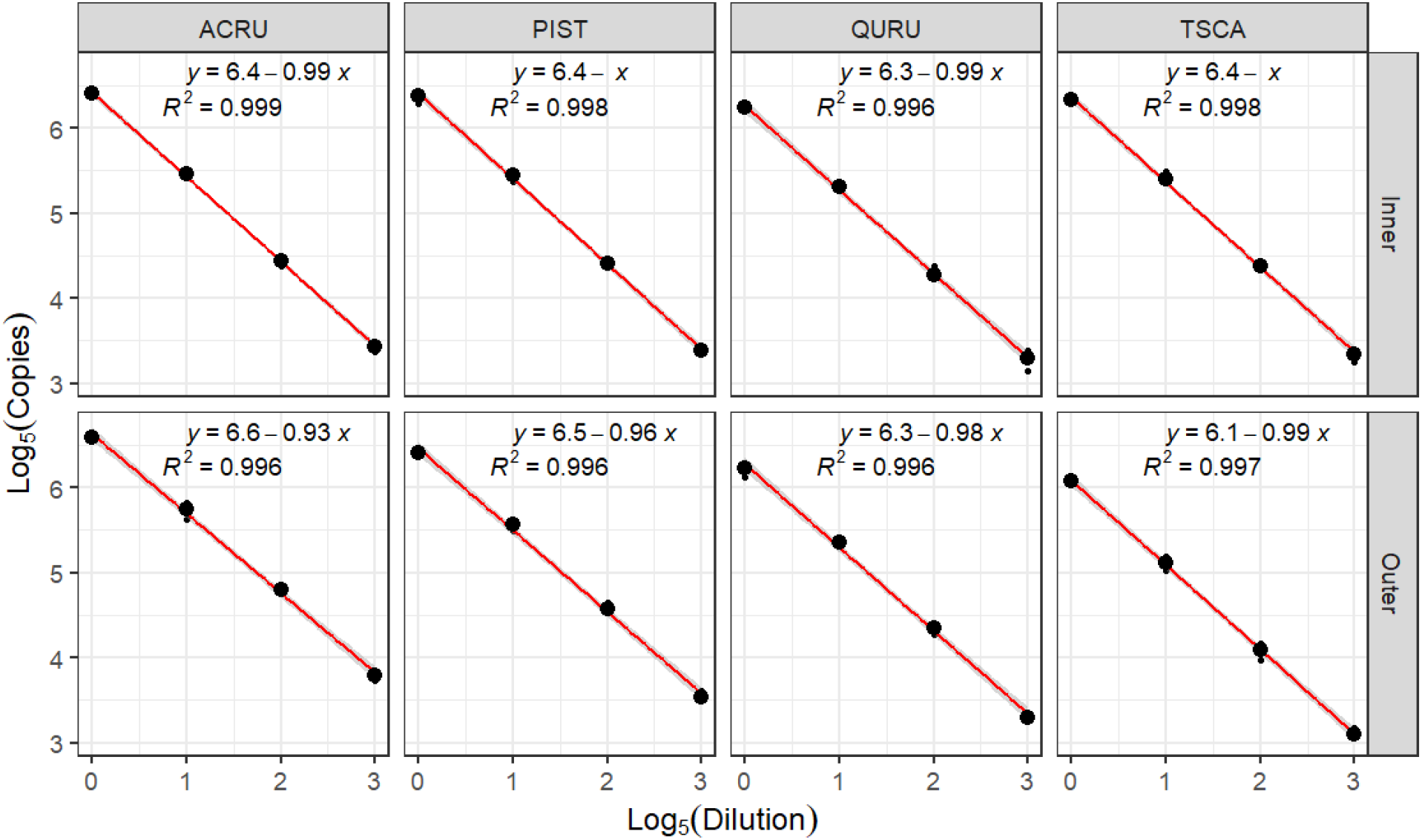
Five-fold dilution (1:1, 1:5, :1:25, 1:125) series, in triplicate, produced using mcrA gBlock-spiked, field-collected samples from January 2023. All samples were extracted using the MagMax Microbiome Ultra Kit, with quantification via the mcrA probe-based ddPCR assay. Samples split by species and tissue-type (inner, i.e. heartwood, and outer, i.e. sapwood), with linear model characteristics displayed in the upper-right. Sample dilution follows linear-behavior, suggesting little inhibition for any of the sample types tested. Each column represents a different tree species: (ACRU: *Acer rubrum*, QURU: *Quercus rubra*, PIST: *Pinus strobus*, TSCA: *Tsuga canadensis*).

### Extraction Efficiency by Substrate

We found that recovery efficiency, defined as the percent recovery of the spiked targets (gBlocks), varied modestly—but significantly—across species for both heartwood and sapwood samples. For 6 of the 8 sample-types tested (4 different species, heartwood/sapwood), average recovery efficiency across the three gene targets (22.2%±3.59%) was significantly lower than the average liquid control (gBlocks spiked into extraction buffer) recovery efficiency (30.4%±3.90%) (**Figure 5**). Red maple sapwood had significantly higher recovery (38%±1.29%) than the control (Paired t-test, p<0.01), while the recovery efficiency for red oak sapwood proved comparable to the control (Paired t-test, p=0.23). In general, we observed that certain woods such as oak and maple ground more easily, and to a finer powder than others, such as hemlock sapwood, which displayed the lowest recovery. For the hemlock sapwood, some larger particles remained after the cryo-grinding process, which may have been resistant to releasing sorbed nucleic acids during the extraction, and thus responsible for the lower recovery. Unspiked, paired controls confirmed that none of the samples had preexisting levels of the target sequences capable of confounding spiking results (**SI Figure 3**).

**Figure 5:**
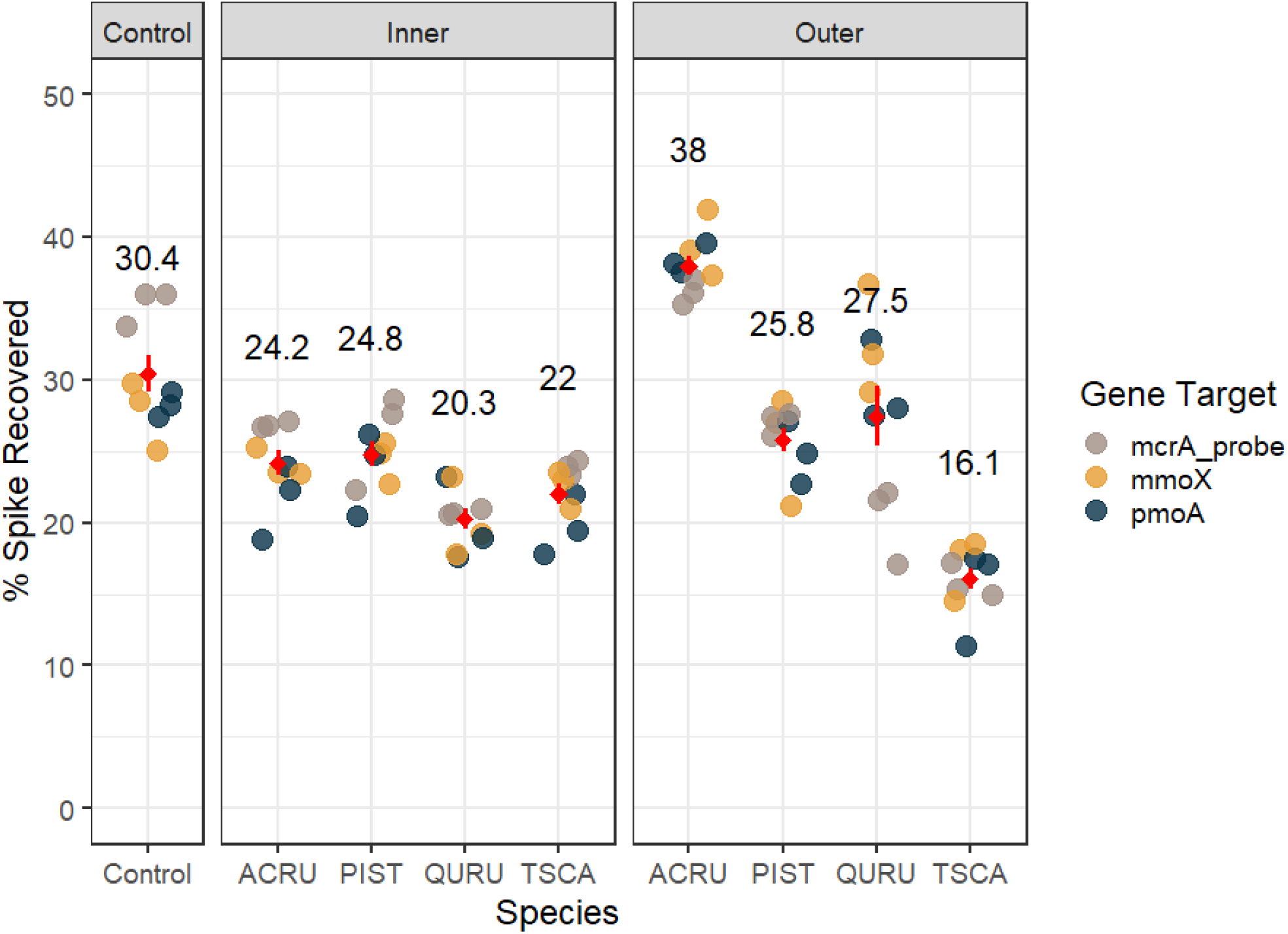
Recovery of gBlock spiking material from January 2023 field-collected samples, as determined by ddPCR. Triplicate extractions, across 4 species and 2 tissues, were performed, along with baseline extractions of gBlocks added directly to extraction buffer. The average recovery across all three triplicates and gene targets is shown with a dot (mean, line representing one SE) and reported above. Each x-axis tick mark label represents a tree species (ACRU: *Acer rubrum*, QURU: *Quercus rubra*, PIST: *Pinus strobus*, TSCA: *Tsuga canadensis*), and “Control” refers to extraction of gBlocks spiked only into extraction buffer, which corresponds to a best-case recovery scenario.

These results suggest that, while there was some species-level variance, recovery efficiencies were generally high, averaging 81.7% that of the idealized control. The lack of clear differences in recovery and PCR-inhibition between angiosperms and gymnosperms, despite contrasting cellular chemistry and structure (Goldstein, 1977), suggests the method is robust, and especially so for heartwood samples. Given our noted challenges with grinding hemlock sapwood, we hypothesize that variations in recovery across species may manifest as a result of indirect impacts on grinding. For instance, the abundance of longitudinal tracheids in hemlock sapwood (Rayirath and Avramidis, 2008) may present issues for grinding, which could be partially ameliorated in tracheid-containing pine species by the presence of potentially weakening structures, like hollow resin canals (Shmulsky and Jones, 2019). Many other factors, like resin content, could further affect grinding efficacy. Should species variance in recovery be due to the indirect impacts of wood structure on processing steps, future optimizations could seek to develop customized grinding protocols for each species (e.g. grinding hemlock samples longer).

### Microbiome Validation

*S*piked cores were used to create surrogate field samples for the controlled, quantitative analysis of method performance given the constraints of field samples, which display natural variability. While a commonly used technique, spiking could produce results that either over-or underestimate the true challenges of extracting DNA from wood endophytes, as we have limited understanding of how endophytic taxa are distributed at a cellular level within wood. To validate our method’s ability to recover environmental endophytic DNA from wood, and across a wider range of taxa, amplicon sequencing was performed using DNA extracted from the heartwood of three additional *Acer rubrum* samples.

Sequencing revealed that DNA from a diversity of taxa (>400 resolved prokaryotic families) were successfully recovered from the three heartwood samples, with similar community profiles observed across all three trees (**Figure 6a**). Moreover, methanogenic taxa, in the families Methanobacteriaceae and Methanomassiliicoccaceae, were found to comprise from 0.48% to 9.9% of all resolved taxa in the heartwood samples (**Figure 6b**). These results demonstrate the method’s ability to recover microbial DNA across a diversity of microbial taxa, including methanogens, our analytical example.

**Figure 6:**
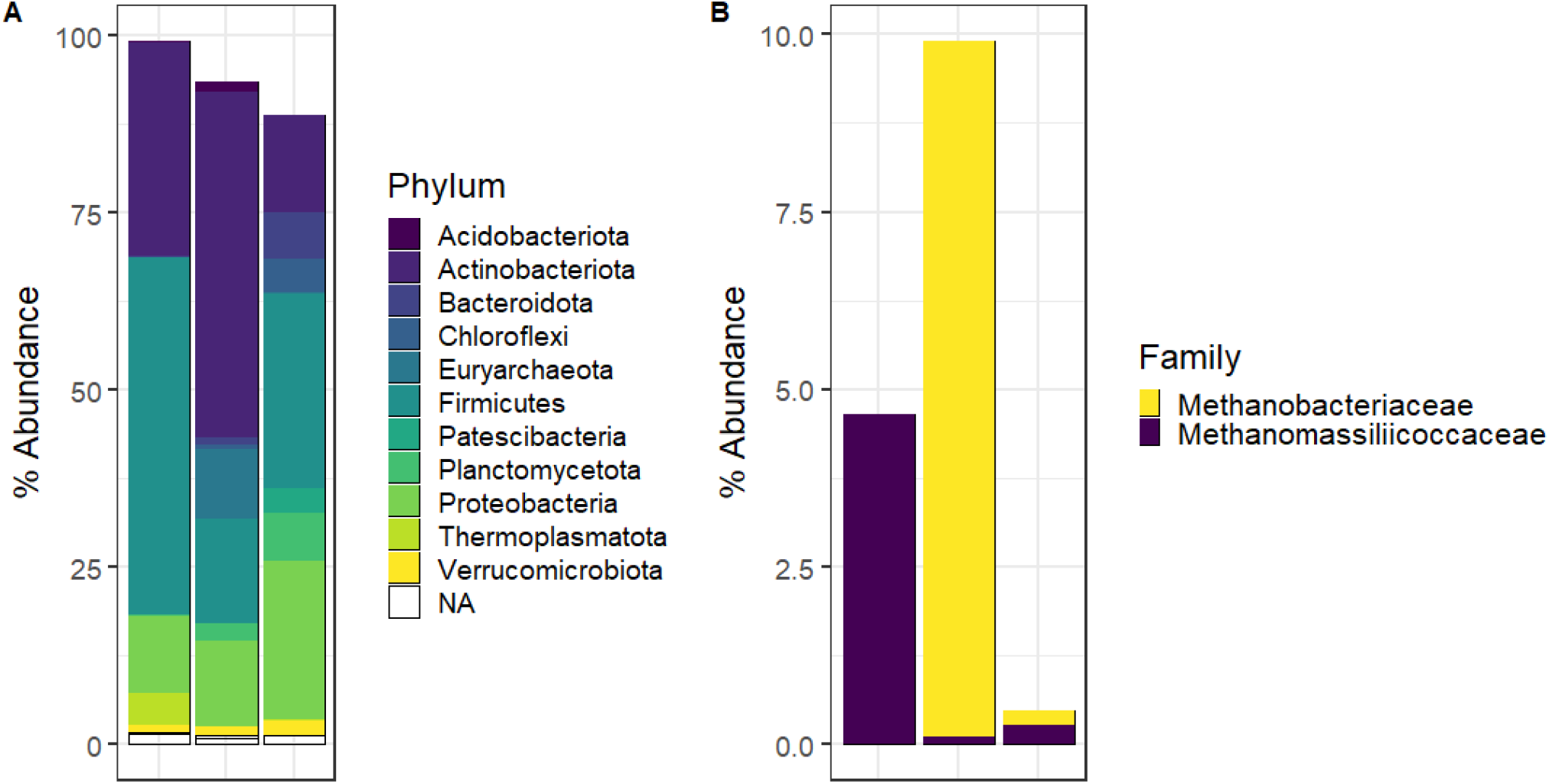
Paired 16S rRNA relative abundance plots produced using microbial DNA extracted from the heartwood of three separate Red Maple (*Acer rubrum*) trees. Even after filtering low abundance taxa **a.)**, many different resolved taxa are recovered from the heartwood samples. When **b.)** filtering taxa for known methanogenic groups, it emerges that methanogenic families make up 0.48% to 9.9% of all resolved taxa in the heartwood samples.

### Losses Throughout Processing

Using sets of sterile dowels, spiked at each step of our methodology (prior to lyophilization, prior to cryogrinding, and prior to nucleic acid extraction), we were able to quantify losses along our processing pipeline. DNA extraction itself contributed the most to spike loss, with freeze-drying and cryo-grinding presenting smaller effects.

Relative to spiked liquid controls, we observed the best recovery for dowel samples spiked at the point of extraction (spike added to ground wood). For samples spiked with culture, we observed a recovery of 34.3% (±10.2) of spiked material, and a 51.7% (±5.2) recovery for samples spiked with gBlocks (**Figure 7**). This recovery, compared to the control, was less than what was observed for spiked field samples (average 81.7%) (**Figure 5**), and was perhaps attributable to imperfect grinding, as the dowels were harder to cryogrind due to the orientation of their grain being parallel to the angle of the grinding impactor (e.g. end-grain cutting board) rather than perpendicular (as was the case with the field-collected cores). This led to a slightly more fibrous powder, which may have been less efficient at releasing spiked material during extractions. Additionally, coarsely ground samples pelleted less effectively during centrifugation steps, leading to greater losses during steps transferring supernatant. This result suggests that sampling living wood parallel to the grain may present more challenge as it relates to homogenization, and that sampling perpendicular to the grain, and across the growth rings, is preferable for grinding.

**Figure 7:**
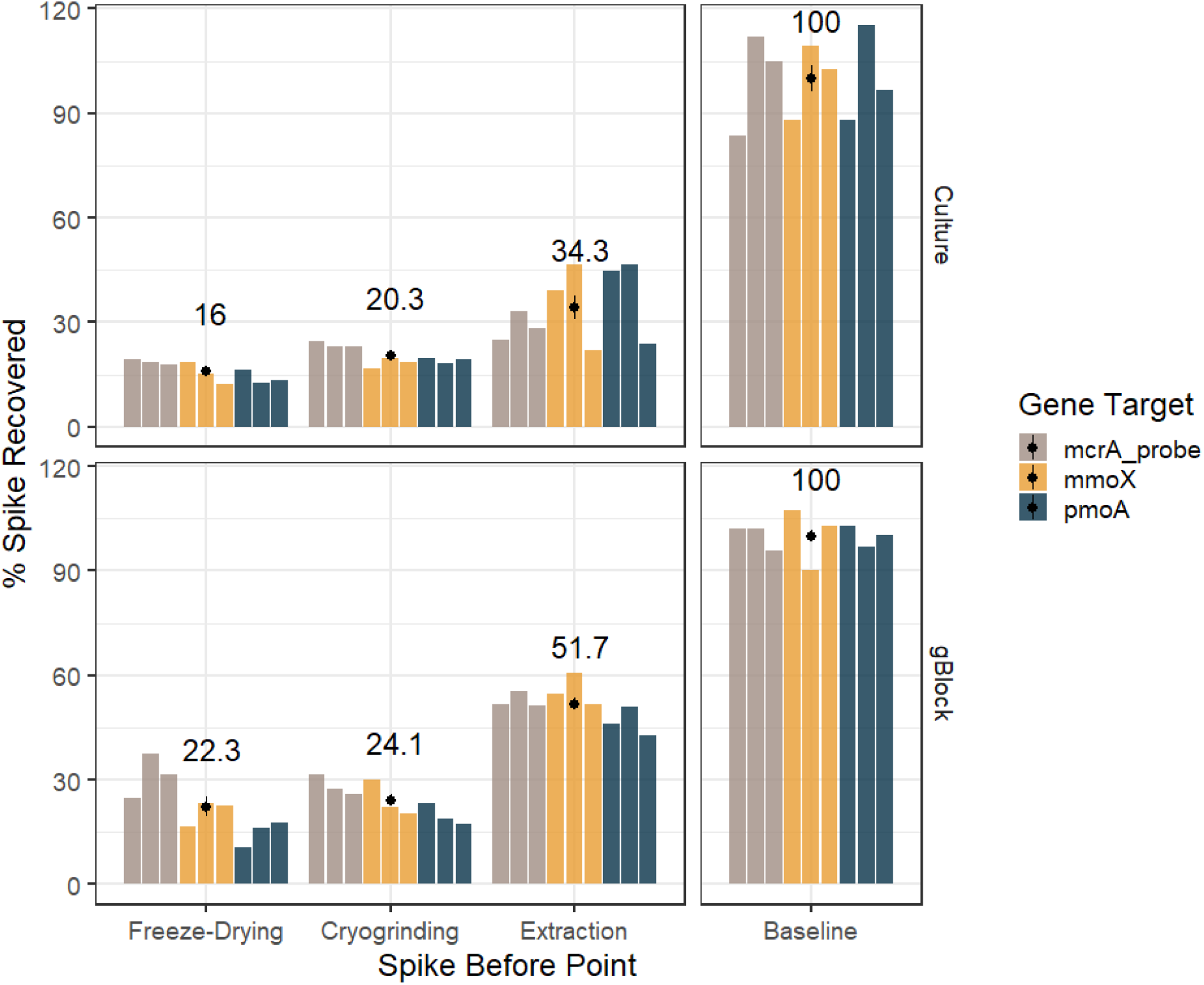
Results of the sequential spiking experiment using sterile dowels, split by spiking material: cultured cells or gBlocks. The mean recovery across all three targets (where each bar is an individual target) is reported above the graph. Recovery percentages were normalized to the baseline kit recovery (ergo average baseline recovery is 100%), which was assessed via extraction of the spiking material when added directly to extraction buffer.

Cryo-grinding was also seen to result in a reduction of recovery efficiency, down to a 24.1% recovery for gBlocks, and 20.3% recovery for culture. Some reduction was, however, anticipated, as material loss (e.g. to the grinding vial) and a modest amount of nucleic acid degradation (shearing) may have occurred during processing. While freeze-drying did not lead to a significant reduction in recovery efficiency for samples spiked with gBlocks, a reduction in spike recovery (down to 16%) was observed due to freeze-drying for culture-spiked samples.

Overall, we observed an additional reduction in spike recovery due to freeze-drying and cryogrinding steps, as recovery, averaged across the two spikes, fell from 43.0% when spiked just prior to extraction to 19.2% when spiked prior to freeze-drying. While this loss is non-trivial, processing wood into a homogenized material is a necessity for consistent and intercomparable nucleic acid extractions of endophytic taxa. Accounting for losses due to the extraction process, losses during handling, and sensitivity of our ddPCR quantification assay, the limit-of-detection (LoD) for our method is between 300-600 cells per 100 mg of sample (dry wood). This LoD is based on 2 μl of eluent used for quantification. Assuming a conservative LoD of 500 cells, this would equate to an abundance of methanogens roughly 200x less than what is typically found in 100 mg of dried soil (Weil *et al*., 2020). As methanogens have been found to be one of the most abundant group within the heartwood of certain tree species (Yip *et al*., 2019; Li *et al*., 2020), and easily detectable via amplicon sequencing, a less sensitive method than either qPCR or ddPCR (Lou *et al*., 2022; Ferreira *et al*., 2023), we are confident in the ability of this method to detect such communities within trees.

Further, this method’s recovery of ∼20%, and LoD of 300 to 600 cells in 100 mg dry wood, fall well within the range of those reported for other environmental samples. In a comparison of six kits for soil extractions, the highest performing kit recovered between 11-35% of spiked material (Dineen *et al*., 2010). Work evaluating extraction of fungal DNA from liquid ranged from 0-30.1% (Fredricks, Smith and Meier, 2005) for a 100 μl sample. Depending on kit and substrate, the LoD for soil samples was observed to be 2-2000 CFU/100 mg dry soil (Whitehouse and Hottel, 2007). Simulated environmental swabs displayed a LoD on the range of 10^3^-10^5^ cells per swab (Dauphin *et al*., 2010), while the extraction of cells collected on a filer was found to have a LoD of 2000-3000 cells per filter (Hospodsky, Yamamoto and Peccia, 2010). As such, the method developed herein performs comparably—or even better—than extraction methods for more common environmental samples, thus opening the study of the endophytic living wood microbiome to the same techniques and scrutiny which have already transformed our understanding of many other ecological niches.

## Conclusions

The method presented herein represents a synthesis of existing techniques, further refined into a high-throughput protocol that allows for the molecular analysis of microbial communities in the living wood of trees and can enable an accelerated study of the tree microbiome. Despite some variance across species, recovery of nucleic acids from most samples was only modestly lower (18.3%) than idealized controls. The LoD of the stated method—500 cells per 100 mg dry wood—is in-line with recovery rates expected for air, water, surface and soil environmental samples, and adequate to detect communities of functional significance. Given that there are over 3 trillion trees on the planet (Crowther *et al*., 2015), even low abundance endophytic taxa could quickly scale-up to a globally significant presence, and thus sensitive detection within the ubiquitous—but poorly characterized—niche of living wood is of global ecological and biogeochemical relevance.

## Acknowledgments

This study was funded by the Yale Center for Natural Carbon Capture (YCNCC) and the Yale Planetary Solutions Project (PSP), and the Yale Institute for Biospheric Studies (to JG). WA was supported by the Department of Defense (DoD) through the National Defense Science & Engineering Graduate (NDSEG) Fellowship Program, and JG was supported through the National Science Foundation (NSF) through the Graduate Research Fellowship Program (GRFP) and a Kohlberg-Donohoe Research Fellowship. We thank Cade Brown, Qespi T’ika Vizcarra Wood, Naomi Norbraten, and Talia Kolodkin for lab and field assistance. We thank Marlyse Duguid and Yale Forests for facilitating this work and Craig Brodersen, Andrew Reinmann, and Clare Kohler for helpful conversations regarding methodology. We thank the Keck Oligo Synthesis Resource at Yale for their assistance with primer production, and the Yale Center for Genetic Analyses of Biodiversity for use of equipment and facilities.

## Author Contributions

WA, JG, and CB developed extraction methods. WA and JG designed and performed all experiments. WA developed and validated ddPCR methods. WA, JG, and JP contributed to the writing of the manuscript. MB, PR, and JP provided revisions to the manuscript and the conception of the work. WA and JG contributed equally to this work.

## Data Availability

All data used in this study are available at: https://datadryad.org/stash/share/X6T_7wvqQmmpnSMh6lhZzeHyFXeA0Q2d3HASqRaPVog

## Supplemental Information

### Probe Selection

Annealing temperatures for both assays were first calculated in-silico using SnapGene, with further refinement via gradient PCR (**SI Table 1**). Annealing temperatures were set as high as possible without observing declines in assay performance. Initial testing of the new primer-probe assays suggested that the set using HEX as a fluorophore was not degenerate enough, as it was capable of detecting mcrA gene fragments derived from *Methanococcus maripaludis*, but not *Methanococcus voltae*. While we increased the degeneracy, and both targets could be resolved, the assay also struggled with separation between positive and negative droplets during ddPCR. For these reasons, the FAM-based set, which presented neither of these issues, was selected over the HEX-based set.

The FAM-based assay displayed strong linearity across a dilution series spanning five orders-of-magnitude, with a slope of 0.99 and an R^2^=0.997. The assay showed consistent detection down to 3 copies/reaction (n=6/6), even averaging 0.42 copies/reaction at a dilution corresponding to 0.33 copies/reaction (n=2/6). As such, the limit-of-detection falls somewhere below 3 copies, and likely is close to a single copy (**SI Figure 2**). Moreover, for all 6 negative controls, the assay produced no false positives.

**SI Table 1:**
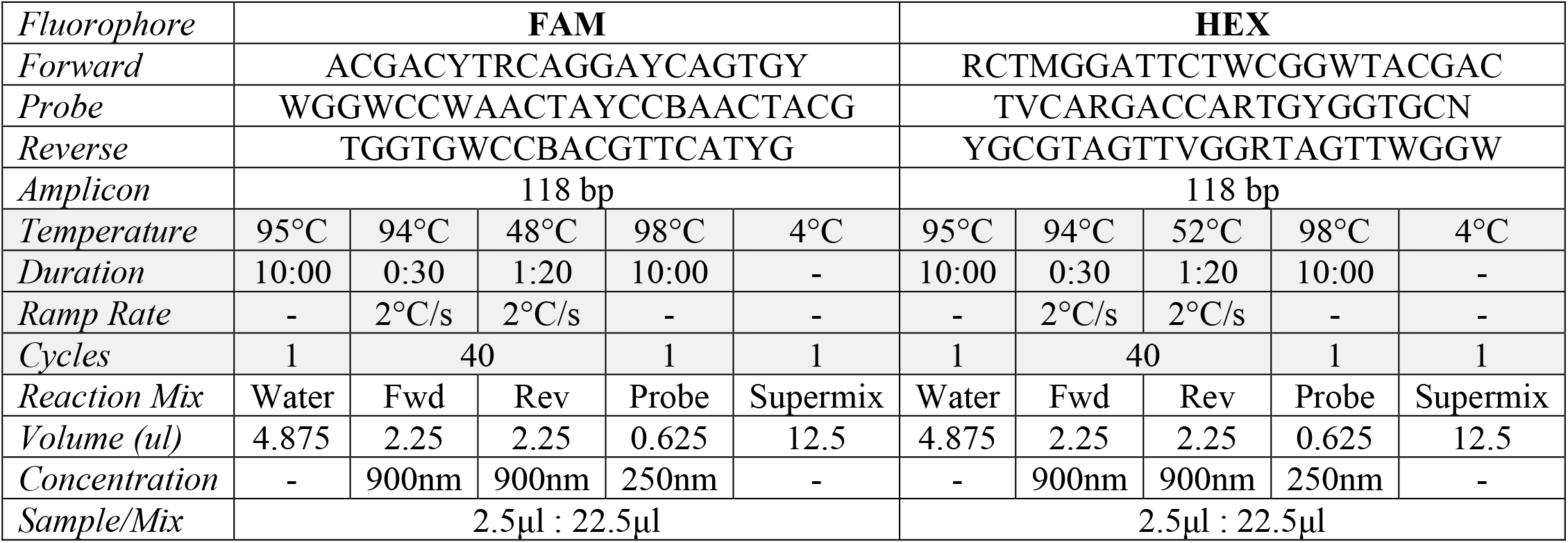
Novel, degenerate ddPCR probes and primers (5’to 3’) targeting the mcrA region, with corresponding thermocycling conditions and reaction mixes. The FAM-based assay was ultimately selected for use in the methods analyses. Both assays used ddPCR Supermix for Probes (No dUTP) (Bio-Rad Laboratories, CA, USA) as the supermix.

**SI Figure 1:**
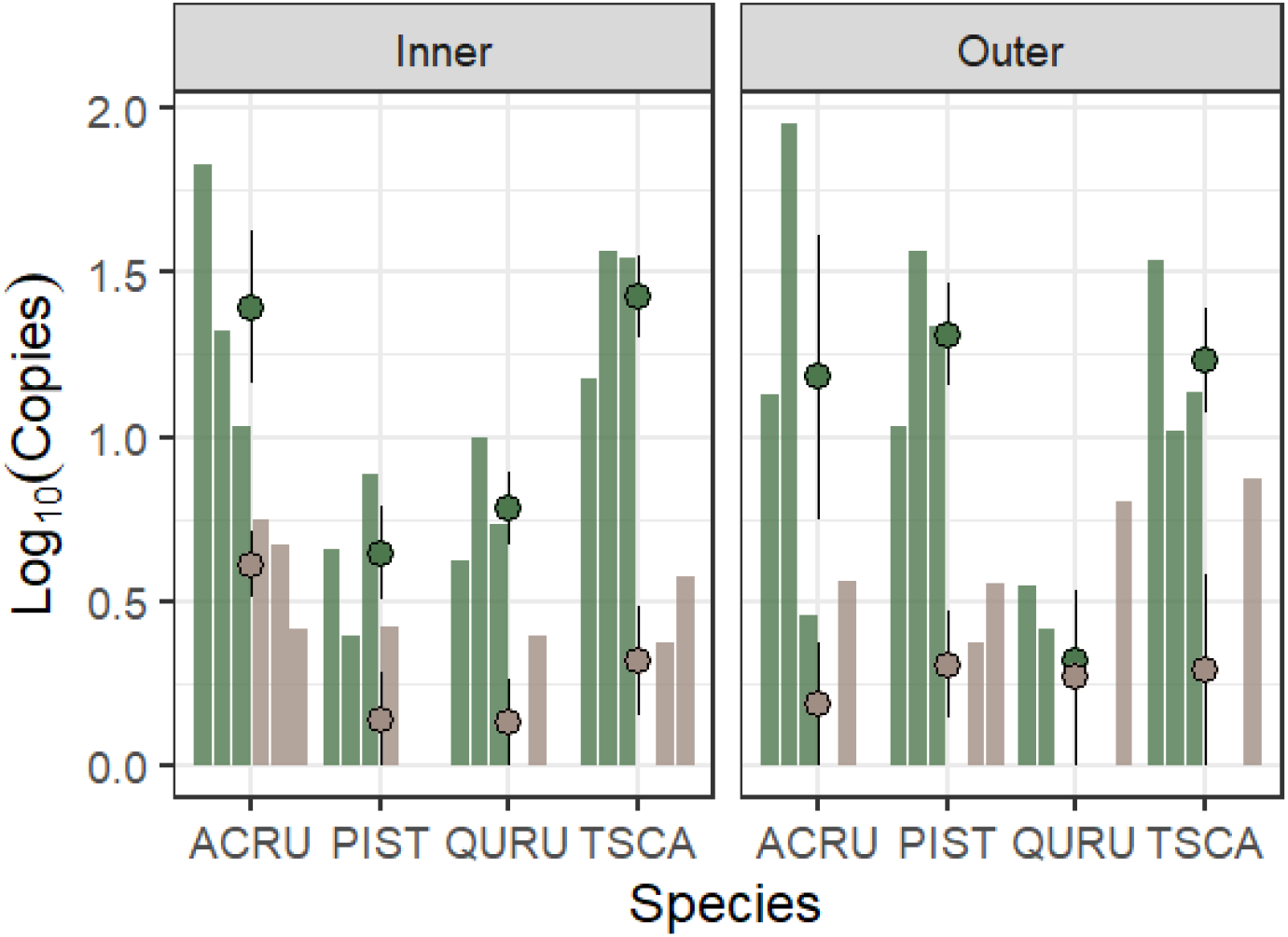
ddPCR using mcrA target assays on unspiked samples, EvaGreen-based assay in green, probe-based assay in tan. Variable rates of false negatives, depending on species and tissue type, were apparent when using existing mcrA primers and EvaGreen chemistry. Bars represent individual samples, and points (with lines) represent the mean (±SE). Samples clustered by species (ACRU: *Acer rubrum*, QURU: *Quercus rubra*, PIST: *Pinus strobus*, TSCA: *Tsuga canadensis*).

**SI Figure 2:**
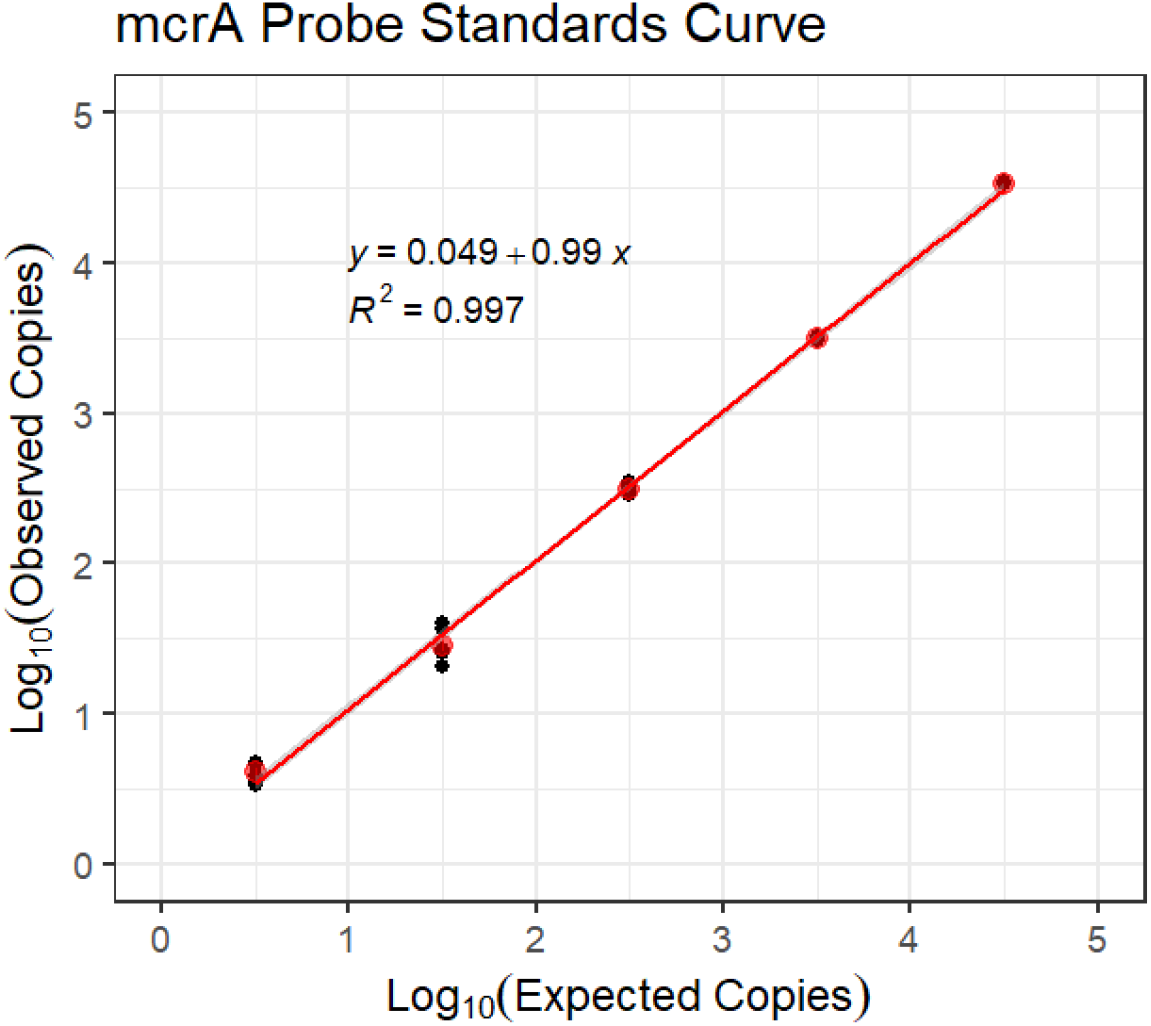
Performance of degenerate FAM-labelled ddPCR assay targeting the mcrA encoding region across five orders of magnitude (n=6 per point). Characteristics of a linear regression fit through the data displayed above.

**SI Figure 3:**
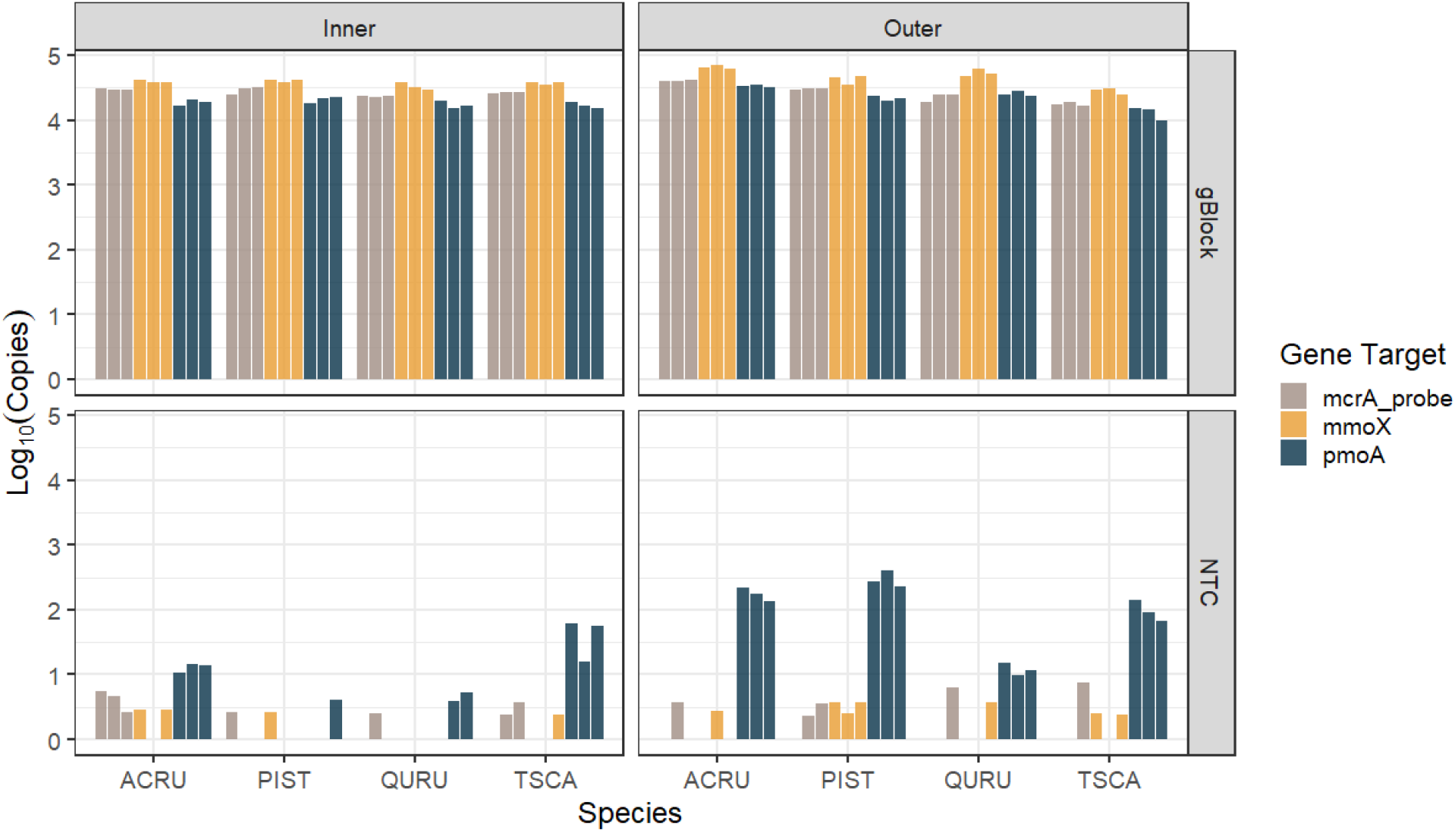
Raw (log10) abundances from the ddPCR performed on January 2023 samples. Spiked samples are shown above, with corresponding no template controls displayed below. While many samples had low abundances of naturally occurring sequences, no sample-type had an amount that would meaningfully confound spike recovery. Each x-axis tick mark label represents a tree species: (ACRU: *Acer rubrum*, QURU: *Quercus rubra*, PIST: *Pinus strobus*, TSCA: *Tsuga canadensis*).

